# On variability in local field potentials

**DOI:** 10.1101/2025.03.27.645661

**Authors:** Mohsen Parto-Dezfouli, Elizabeth L. Johnson, Eleni Psarou, Conrado Arturo Bosman, B. Suresh Krishna, Pascal Fries

## Abstract

Neuronal coding and decoding would be compromised if neuronal responses were highly variable. Intriguingly, neuronal spike counts (SCs) show a reduction in across-trial variance (ATV) in response to sensory stimulation, when SC variance is normalized by SC mean, that is, when using the Fano factor ^1^. Inspired by this seminal finding, ATV has also been studied in electroencephalography (EEG) signals, revealing effects of various stimulus and cognitive factors as well as disease states. Here, we empirically show that outside of evoked potentials, the ATV of the EEG or local field potential (LFP) is very highly correlated to the intra-trial variance (ITV), which corresponds to the well-known power metric. We propose that the LFP power, rather than the raw LFP signal, should be considered with regard to putative changes of its variability. We quantify LFP power variability as the standard deviation of the logarithm of the power ratio between an active and a baseline condition, normalized by the mean of that log(power ratio), that is the coefficient of variation (CV) of the log(power ratio). This CV(log(power ratio)) is reduced for gamma and alpha power when they are enhanced by stimulation, and it is enhanced for alpha power when it is reduced by stimulation. This suggests a potential inverse relation between changes in band-limited power and the corresponding CV. We propose that the CV(log(power ratio)) is a useful metric that can be computed for numerous existing and future LFP, EEG or MEG datasets, which will provide insights into those signals’ frequency-specific variability and how they might be used for neuronal coding and decoding.

## Introduction

The presence and the properties of a visual stimulus can be decoded from the spike counts (SCs) of neurons in early visual cortex. These SCs can be considered a signal in the context of signal detection theory (SDT, ^2^). In SDT, the signal emitted by a given neuron in response to a given stimulus is ideally identical for each presentation of that stimulus, i.e., there is no noise or variability across trials. However, empirically, different trials of presentation of a given stimulus lead to SCs that vary, i.e., there is across-trial variability (ATV). ATV of SCs has been found in many brain areas of many species ^1, 3–8^. Intriguingly, ATV of SCs in non-human primate superior temporal sulcus predicts choice behavior during social interaction: Neurons selective for social interactions show reduced ATV, and correct social behavior is predicted by reduced ATV ^9^. ATV has been prominently investigated with regard to variability in neuronal SCs. As SC variance is expected to scale linearly with SC mean for a Poisson or quasi-Poisson process, the ratio of SC variance over SC mean, the Fano Factor (FF), has been used to quantify mean-adjusted SC variability ^1^. The SC FF is reduced by stimulus onset, leading to the conclusion that stimulus onset quenches neuronal variability ^1^. Importantly, this also holds when stimulus onset does not increase firing rates ^1^. In addition to sensory stimuli, non-sensory factors such as arousal, attention, and adaptation can also reduce neuronal variability ^7, 10^.

More recently, ATV has been investigated for continuous signals, i.e., EEG or magnetoencephalography (MEG), with the intention to investigate whether stimuli or cognitive tasks similarly quench neuronal variability (for review see ^11^). While studies on ATV in SCs have typically used the FF and thereby normalized SC variance by SC mean, later studies on ATV in continuous signals have typically used the raw ATV without normalization by the mean. Those studies found e.g. that the ATV of EEG signals recorded over task-relevant brain areas is reduced by spatial attention ^12^ and decision making ^13^. Moreover, ATV is altered during cognitive impairments. For instance, patients with autism, attention deficit hyperactivity disorder, and schizophrenia exhibit increased ATV, which has been interpreted as indicating greater instability of the respective neuronal systems due to these conditions ^14–18^.

Here, we calculated ATV of the LFP recorded from macaque electrocorticography (ECoG) arrays and found that attention reduced ATV in the alpha-beta band while it increased ATV in the gamma band, similar to changes in LFP power in the corresponding frequency bands ^19^. We further observed that the ATV of the ongoing LFP outside of event-related potentials (ERPs) was highly correlated to intra-trial variability (ITV). Importantly, ITV corresponds to the LFP power, as studied in numerous LFP, EEG and MEG studies. During ERPs, ATV is reduced compared to ITV, and this reduction is correlated to the degree that the ERP explains total power. For human EEG data, we observed highly similar results. We demonstrate by simulation that for ongoing activity, ATV and ITV are expected to be identical under various conditions related to the equivalence of time and ensemble averages (ergodicity). We then show how this equivalence is broken and ATV can be lower than ITV via the addition of a constant component across trials ^20^. Finally, we constructed a metric that meaningfully captures the variability for LFP power, as opposed to the raw LFP waveform, and that is mean adjusted, like the FF, in order to de-emphasize power-variance changes that are simply due to changes in mean power. As LFP power is log-normally distributed, we constructed this metric as considered the coefficient of variation (CV) of the log10-transformed LFP power ratio between an active and a baseline condition. This metric revealed intriguing effects. When visual stimulation increased visual cortical LFP gamma power, it reduced the CV(log(power ratio)) in the gamma band. For alpha power, the effect depended on stimulus-induced changes: In one dataset with intracortical microelectrode recordings, stimulation increased alpha power and reduced its CV; in another dataset with ECoG recordings, stimulation decreased alpha power and increased its CV. Together, these results suggest that the CV(log(power ratio)) is a meaningful metric of the mean-adjusted variability of LFP power, and that this metric is inversely related to changes in power. These findings are relevant for the use of different LFP frequency bands for decoding brain activity.

## Results

### ATV resembles power

First, we calculated ATV of the LFP recorded from an ECoG grid covering 15 brain areas of an awake macaque monkey. During the experiment, the monkey kept its gaze on the fixation point while performing an attention task. The attention task resulted in two conditions (see Methods for details): (1) a condition with attention to the hemifield contralateral to the recorded brain areas, referred to as the attend IN condition (as most visually responsive brain areas are more sensitive for stimuli in their contralateral hemifield), (2) a condition with attention to the ipsilateral hemifield, referred to as attend OUT condition. Figure 1a shows LFP traces of 150 example trials from one electrode on area V1 during fixation and stimulus presentation. In line with the literature, ATV of wideband LFP decreased following stimulus onset (Figure 1d). Subsequently, we explored ATV of LFP data filtered in the alpha-beta and the gamma bands. To this end, LFPs of the example trials were bandpass filtered in the alpha-beta band (8-18 Hz) and the gamma band (70-80 Hz). For each band, ATV was calculated and investigated as a function of time around stimulus onset. Stimulus onset led to a decrease of ATV in the alpha-beta band along with an increase in the gamma band (Figure 1b,c,e,f). These ATV changes were consistent with modulations of LFP power for the corresponding frequency bands (Figure 1g).

**Figure 1.**
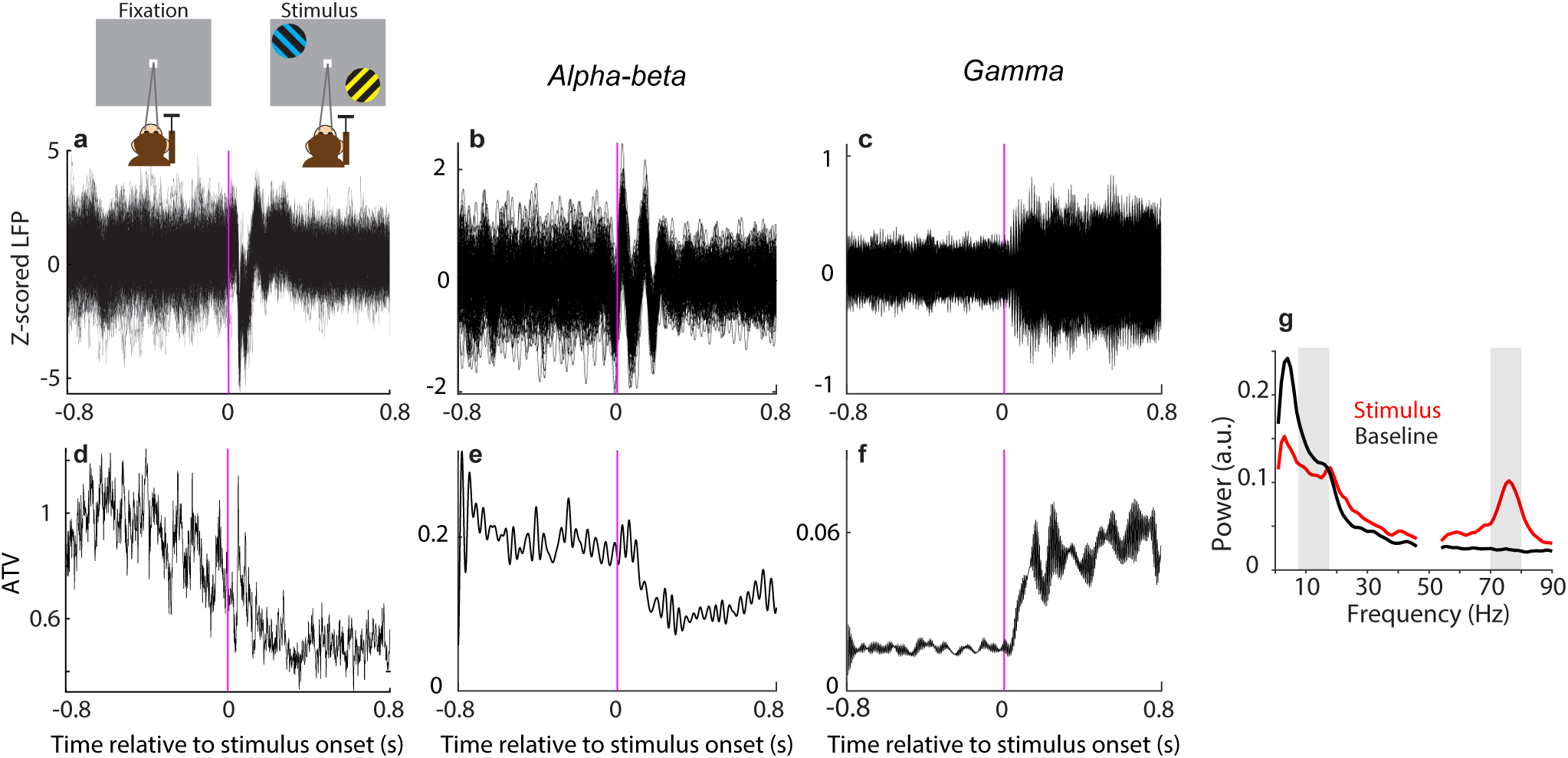
Across-trial variability (ATV). **a**) LFP from 150 sample trials recorded from a single channel in area V1 during fixation and stimulus presentation. **b, c**) Same data as (**a**), but after filtering in the alpha-beta frequency band (8-18 Hz; **b**) or in the gamma frequency band (70-80 Hz; c). **d-f**) Across-trial variance (ATV), during fixation and stimulus presentation for wideband signal (**d**), alpha-beta-band signal (**e**) and gamma-band signal (**f**). **g**) Comparison of power spectra between stimulus and baseline periods. Shaded bars indicate data in alpha-beta and gamma bands.

To further investigate the relationship between power and ATV, we focused on two clusters of areas based on their dominant peak power, as we have established previously ^21^: an occipital cluster containing areas V1, V2 and V4, and a frontal cluster containing areas F1, F2 and F4. In these two clusters of areas, we calculated ATV in the alpha-beta and gamma bands, respectively. ATV was computed for each time point across four periods of the task: fixation (last 0.8 s before stimulus onset), stimulus onset (0.8 s following stimulus onset), attention-cue onset (0.8 s following cue onset), and sustained attention (last 0.6 s before stimulus change). ATV in the alpha-beta band showed decreases after the stimulus onset and after the cue onset (Figure 2a). The decrease after the cue onset appeared stronger for attend-IN versus attend-OUT, in particular for the frontal areas (Figure 2a red versus black lines). ATV in the gamma band showed a first increase after stimulus onset and a further increase after cue onset (Figure 2b). In the occipital areas, the increase after cue onset appeared stronger for the attend-IN than the attend-OUT condition, and this difference was further accentuated during the last 0.6 s before the stimulus change (Figure 2b red versus black lines). Intriguingly, the corresponding power spectra reveal similar reductions of the alpha-beta power (Figure 2c) and increases of the gamma power (Figure 2d) for the attend-IN compared to the attend-OUT condition. We refrain from presenting statistics for these effects here, as they (or closely related effects from the same dataset) have been presented previously (Richter, Thompson et al. 2017).

**Figure 2.**
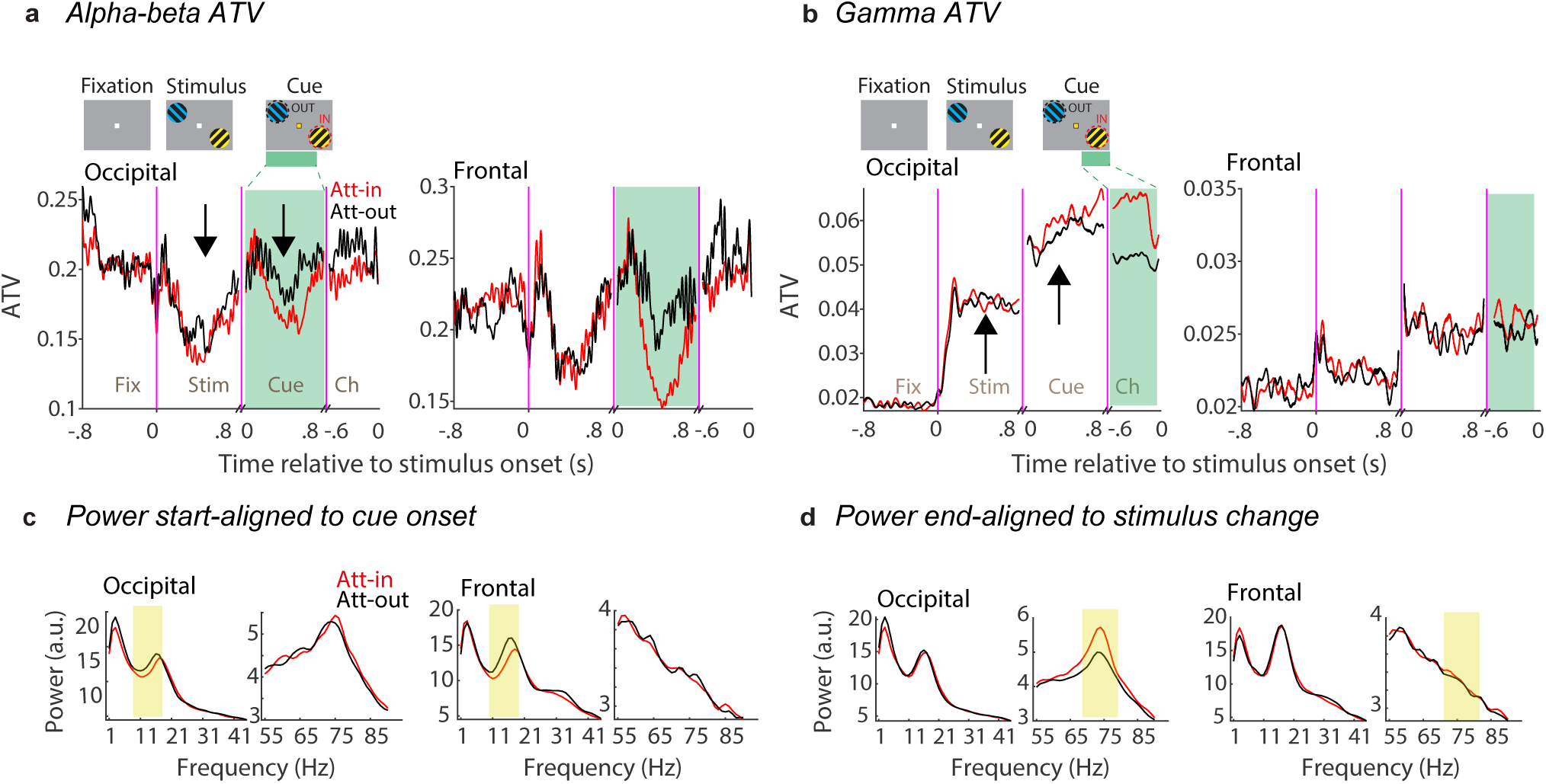
ATV and power spectra. **a**) Time-course of average ATV of the LFP filtered in the alpha-beta band (8-18 Hz) over occipital (left) and frontal (right) clusters for attend-IN (red) and attend-OUT (black) conditions. Arrows indicate the reduction of ATV during stimulus presentation and attention periods. Results were smoothed using a 20 ms sliding window. *Fix*: fixation, *Stim*: stimulus, *Cue*: aligned to attention cue onset, *Ch*: aligned to stimulus change. **b**) Same as (**a**), but for the LFP filtered in the gamma band (70-80 Hz). Green shaded areas were used for power analysis. **c, d**) LFP power spectra for attend-IN (red) vs. attend-OUT (black) conditions based on 0.3-0.8 s after cue onset (c), and based on 0.5 s before the stimulus change (**d**). Yellow shaded areas indicate frequency ranges of alpha-beta and of gamma used in (**a, b**).

### ATV is highly correlated to ITV, which corresponds to power

These results suggest that ATV might be related to power. Whereas, ATV is calculated across trials per time point, power is calculated across time points per trial, typically followed by averaging over trials. A proxy for power can be calculated by rectifying the signal, which is the absolute value of the signal, before averaging over trials. This average rectified LFP showed stimulus-, cue-, and attention-related dynamics in the alpha-beta band (Figure S1a,b) and gamma band (Figure S1c,d) that were similar though not identical to the ATV (Figure 2a,b).

We further investigated this putative relation between ATV and power. The power of a signal corresponds to the variance of the signal across time, defined as the sum of the squared deviations from the signal mean. Thus, signal power corresponds to the ITV, and we use this acronym, because it exposes the similarities and differences between ATV and ITV: ATV quantifies the variability across trials per time point, and can subsequently be averaged over time points; ITV quantifies the variability across time points per trial, and can subsequently be averaged over trials. If the signal has been filtered, ITV corresponds to the power in the corresponding frequency band; if the signal has not been filtered, ITV corresponds to the total power of the signal. For direct comparison between ITV and ATV, we opted to use the unfiltered signal, because it has been used for most applications of ATV on continuous neuronal signals in the literature. While ATV has typically been calculated separately per time point, this is not possible for ITV. Rather, the calculation of ITV, both for the unfiltered or the filtered signal, requires multiple samples per trial and thereby some time period. Therefore, to directly compare ITV and ATV for the same data, we defined three periods: (1) the fixation period (last 0.5 s before stimulus onset), (2) the event-related potential (ERP) period (first 0.3 s following stimulus onset), and (3) the sustained-response period (0.3 to 0.8 s after stimulus onset). ITV was calculated per trial and per period across the time points, and then averaged over trials. ATV was calculated across trials per time point, and then averaged over time points (see Figure 3a for a schematic illustration).

**Figure 3.**
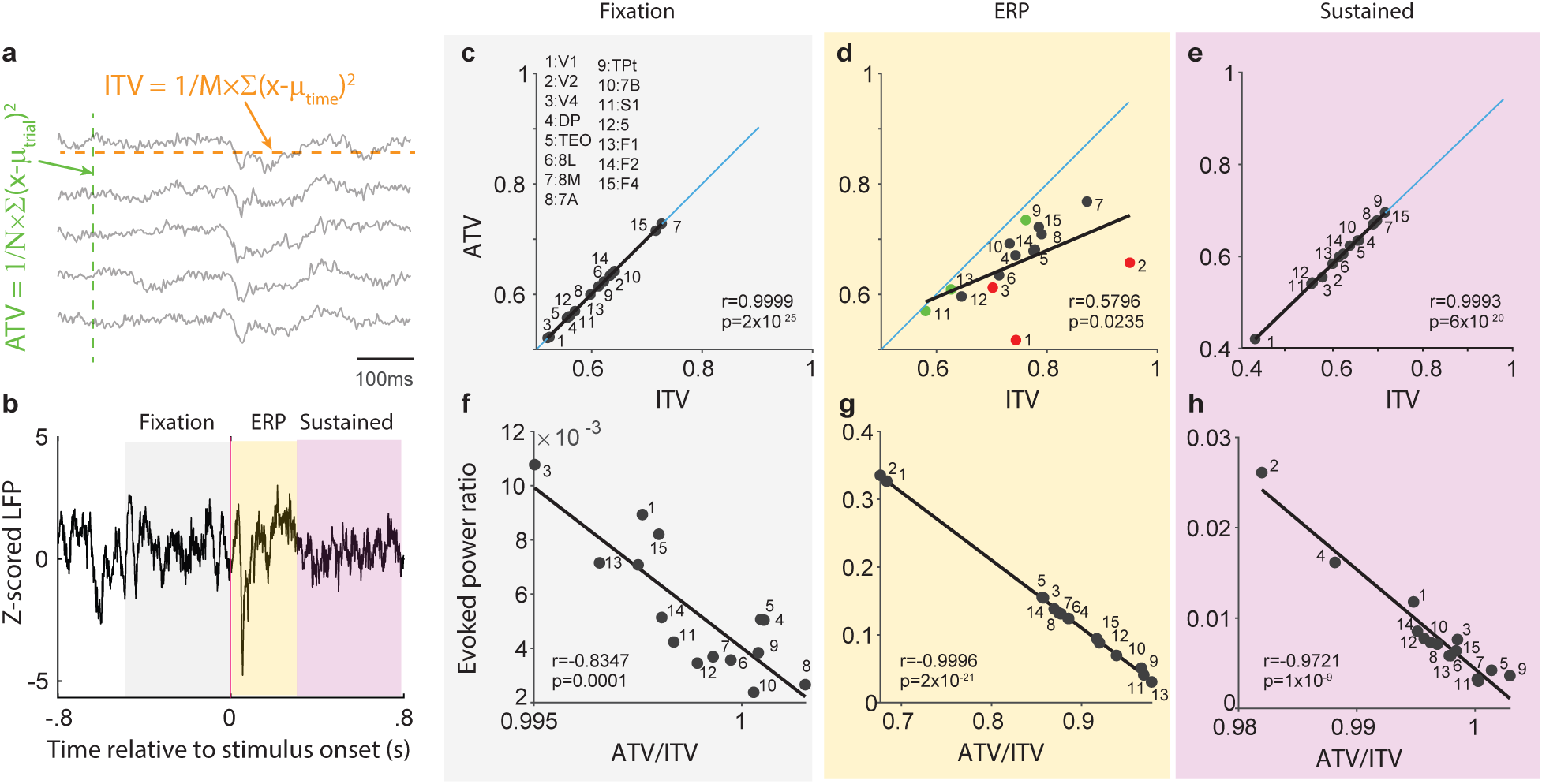
ATV correlates with ITV. **a**) Schematic illustrating the estimation of ATV and ITV for five sample trials. ATV is first calculated per time point, as the variance across trials and subsequently averaged over time points (this last step is not illustrated here). ITV is first calculated per trial, as the variance across time points within the trial, and subsequently averaged over trials (this last step is not illustrated here). **b**) Averaged z-scored LFP for three task periods: Fixation (gray; −0.5-0 s relative to stimulus onset), ERP (yellow; 0-0.3 s relative to stimulus onset) and sustained-activity period (purple; 0.3-0.8 s relative to stimulus onset). **c**) Scatter plots of ATV versus ITV across the 15 indicated brain areas for the Fixation period as indicated by the corresponding gray shading between panels **b** and **c**. The lower right corner provides the Pearson correlation coefficient between ATV and ITV across the 15 areas. The blue diagonal line indicates the line of equality between ATV and ITV. **d**) Same as (**c**), but for the ERP period. **e**) Same as (**c**), but for the sustained activity period. **f-h**) Correlation between the evoked power ratio, that is, the ratio of the evoked power over the total power, and the ratio of ATV over ITV, separately for each task period.

For the direct comparison between ATV and ITV, we used the same ECoG data as reported above. Both ITV and ATV were calculated for individual recording channels and subsequently averaged over channels within each recording area, separately for the three periods as indicated by the colored backgrounds in Figure 3b. Figure 3c-e shows scatter plots of ATV versus ITV across 15 brain areas, separately for the aforementioned periods of fixation (Figure 3c), ERP (Figure 3d), and sustained-response (Figure 3e). Each scatter plot shows the diagonal as a blue line, with points on/above/below the diagonal showing that ATV is equal to/larger/smaller than ITV. In addition, each plot shows the linear regression of ATV versus ITV as black line. This reveals a significant positive correlation between ATV and ITV in all three periods. Notably, the regression coefficient between ATV and ITV assumes values of 0.9999 (Figure 3c) and 0.9993 (Figure 3e) during the fixation and sustained-response periods, respectively, with all points lying essentially on the diagonal, suggesting that ATV and ITV across those 15 brain areas are essentially identical. Intriguingly, during the ERP (Figure 3d), ATV for many brain areas is reduced compared to ITV.

The amount of this reduction varies across areas, such that the regression coefficient between ATV and ITV across areas is reduced, yet still significant (r=0.5796, p=0.0235).

### ATV is reduced below ITV when signal power is phase aligned to an event

A closer inspection of the data points for individual areas reveals that the ATV reduction is particularly strong for early and intermediate-level visual areas V1, V2 and V4 (red dots in Figure 3d), known to show relatively strong ERPs to visual stimulus onset. By contrast, the ATV reduction is comparably weak for areas Tpt, S1, and F1 (green dots in Figure 3d), which are not expected to show strong ERPs for visual stimuli. Thus, ATV and ITV are almost identical in the absence of ERPs, whereas during ERPs, ATV is reduced compared to ITV, and this reduction is likely related to the ERP strength.

To further investigate the relationship between ATV reduction and ERP strength, we assessed ERP strength by calculating the ratio of evoked power to total power. For the calculation of total power, power is calculated per trial and subsequently averaged over trials. By contrast, for the calculation of evoked power, the LFP is averaged over trials, resulting in the ERP, and subsequently, the power of this ERP is calculated. Evoked power can range between zero and the value of total power, such that the ratio of evoked over total power assesses the fraction of total power that is due to the ERP; we refer to this as the “evoked power ratio” (y-axis values in Figure 3f-h). As expected, the evoked power ratio was large during the ERP period, ranging from 0.025 for areas Tpt, S1 and F1, to >0.3 for areas V1 and V2 (Figure 3g). As also expected, the evoked power ratio was substantially smaller during the sustained response period (Figure 3h), and even smaller during the pre-stimulus baseline (Figure 3f). Importantly, the evoked power ratio was negatively correlated to the ratio of ATV over ITV for all three task periods. For the ERP period, this correlation assumed a value of 0.9996, suggesting that the ATV reduction is most likely due to the ERP (Figure 3g).

Given that most ATV studies have used human data, we extended our analysis to test the relationship between ATV and ITV using 64-electrode EEG data from one human subject ^22^. Both ATV and ITV were computed during four distinct periods of a working memory task (Figure S2a; see Methods for details). The time course of the ATV averaged over the 64 EEG electrodes revealed a noticeable decrease during stimulus presentation and working memory retention periods (Figure S2b). For each of the four task periods, and each of the 64 electrodes, ATV and ITV were calculated. Within each task period, the correlation between ATV and ITV was assessed across 64 electrodes (Figure S2c). The results consistently showed very strong correlations (r≈0.99), suggesting that ATV and ITV are essentially identical for human EEG data. Note that in these data, we did not observe clear reductions of ATV after an external event, most likely reflecting weak phase resetting in this dataset, because stimuli varied across trials in their location, shape and color.

### Empirical relationships between ATV, ITV and ERPs can be simulated very easily

The above results in macaque ECoG and human EEG data suggest that ATV is almost identical to ITV for periods without ERPs, whereas ATV is reduced relative to ITV for periods with ERPs. To further investigate this, we used a few very simple simulations. First, we simulated 100 trials of normally distributed white noise, whose amplitude was either decreased (Figure 4a) or increased (Figure 4b) at time zero. These changes in white-noise amplitude constitute changes in broadband power, i.e., broadband ITV. As ITV was essentially identical to ATV in experimental data without ERPs, we expected the same to hold for those simulated data. Indeed, Figure 4d,e shows that a decrease or increase in intra-trial amplitude led to a decrease or increase in cross-trial ATV, respectively. We simulated 20 electrodes, each of which with an individual level of power, generated by an individual SD of the simulated white-noise process. Across the 20 electrodes, the regression coefficient between ATV and ITV assumed the value of one, that is, ATV and ITV were identical (Figures 4g,h). These simulation results were expected from a white-noise process, where each data point is drawn independently, such that variance calculated across time or trials is expected to be the same, and we show this mainly for illustration and as context for the next simple simulation.

**Figure 4.**
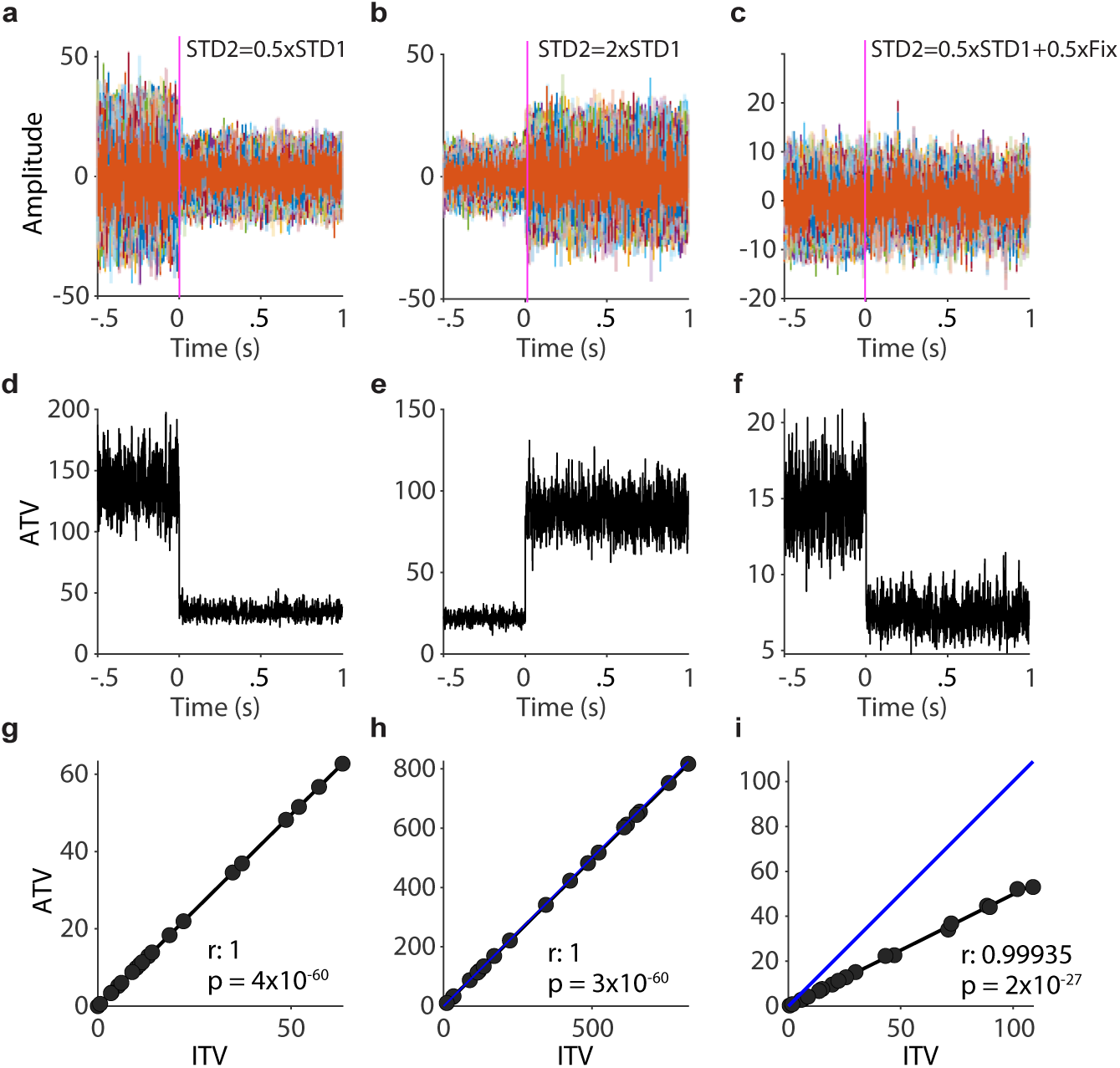
ATV change is driven by amplitude modulation or across-trial phase alignment. **a**) Sample of 100 simulated trials drawn from two distributions, representing a fixation period (−0.5-0 s) and a stimulus period (0-1 s). Both distributions follow a normal distribution defined as 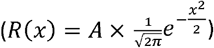. The stimulus period has half the STD compared to the fixation period. **b**) Same as (**a**), but the STD of stimulus period was doubled compared to the fixation period. **c**) Illustration of the effect of phase alignment for ATV and ITV. The signal before time zero was generated as the superposition of two independent white-noise processes of equal mean and SD, which were also independent across trials. The signal after time zero was generated equally, except that one of the two white-noise processes was constant across trials. **d-f**) ATV corresponding to a (**d**), **b** (**e**), and **c** (**f**). **g-i**) Correlation between ATV and ITV corresponding to (**a-c**), respectively. Each dot represents data from one of 20 assumed channels. The blue line indicates equality between ATV and ITV.

Next, we simulated an ERP that was generated by a partial phase reset at time zero ^23^. A phase reset leads to a signal component that is event-locked across trials. We simulated this event-locked component as a white-noise process that was identical for all 100 trials. This event-locked component was not supposed to change the overall power, i.e., ITV of the signal, which was accomplished as follows: The signal before time zero was generated as the superposition of two independent white-noise processes of equal mean and SD, which were also independent across trials. The signal after time zero was generated in the same way, except that one of the two white-noise processes was constant across trials. The results are shown in Figure 4c,f,i: At time zero, the raw signals across the 100 trials show no apparent change (Figure 4c), yet the ATV was strongly reduced (Figure 4f). Across the simulated electrodes, the regression coefficient between ATV and ITV was 0.99, yet the regression slope was reduced to 0.496, that is, to a value very close to half (Figure 4i). This illustrates how a constant component across trials, accounting for half of the total signal energy, in the absence of changes in power, i.e. ITV, can lead to reductions in ATV.

### A mean-adjusted metric for LFP variability

The investigation of ATV in continuous data has often been motivated by the prominent finding that stimulus onset or other task-relevant events, quench neuronal variability ^1^. This variability is first measured as single-unit SC variance. Subsequently, to minimize confounding effects of changes in SC mean, the SC variance is normalized by the SC mean, which corresponds to the SC FF. We calculated the single-unit SC FF in intracortical microelectrode recordings from awake macaque area V1 (Figure 5a), and confirmed that it was reduced after stimulus onset as previously reported (Figure 5b, N=232 single units recorded in 22 sessions in one macaque). We next derived a similar metric for the simultaneously recorded LFP data. Raw LFP amplitude values fluctuate around zero, thereby taking both positive and negative values, and, outside of ERPs, average out to zero but do not tend towards particular non-zero values. By contrast, SCs take only positive values, with values that are specific to the respective neuron and experimental condition, e.g. the respective sensory stimulus. A particular single neuron in visual cortex may on average during a one-second trial show a SC of 5 in the absence of a visual stimulus, and of 50 in the presence of a visual stimulus. Similar properties exist for LFP power: LFP power takes by definition only positive values, with values that are specific to the respective recording site and experimental condition. For example, the LFP recorded from a particular site in visual cortex may show an average gamma-band power of x μV^2^/Hz in the absence of a visual stimulus, and of 5x μV^2^/Hz in the presence of a visual stimulus.

**Figure 5.**
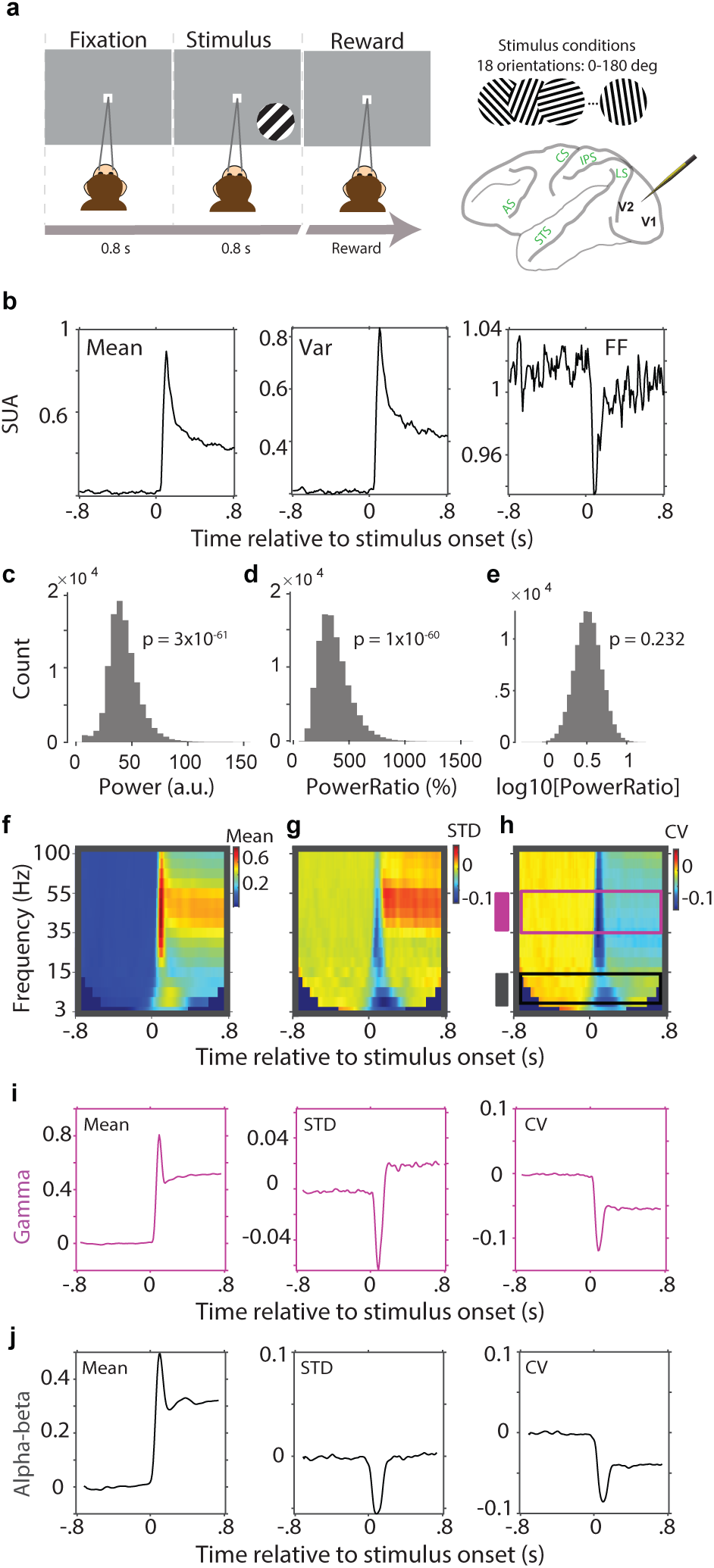
The Coefficient of Variation (CV) of log(power) as a mean-adjusted variance metric for LFP power. Data from intracortical microelectrode recordings including single-unit recordings. **a**) Schematic representation of the passive fixation task during which microelectrode data were recorded. For each session, one out of 18 possible orientations was selected as stimulus and presented for 90 trials. Data were recorded simultaneously from 16 electrodes inserted into areas V1 and V2. **b**) Spike-count mean (left), variance (middle), and Fano Factor (FF; variance divided by mean), computed within 30 ms sliding windows. **c**) Distribution of raw LFP gamma-band power across sessions, channels and trials. **d**) Same as (**c**), but for the ratio of gamma-band power during stimulation over gamma-band power during pre-stimulus baseline. **e**) Same as (**d**), but taking the log10 of the ratio. All subsequent panels use this log10-ratio. **f-h**) Time-frequency representation of the log10-ratio mean (**f**), standard deviation (**g**), and coefficient of variation (**h**). Shown are the averages over 90 repetitions of a single stimulus. **i**) Cross-cuts for the gamma band indicated in (**h**) as a purple rectangle and shown here as time-course to better illustrate the stimulus-related dynamics. **j**) Same as (**i**), but for the alpha band indicated in (**h**) as a black rectangle.

It is an interesting question whether stimulus onsets or other relevant events also quench LFP power variability as they do for single-unit SC. Single-unit spike-trains have often been modeled as Poisson processes, for which the variance equals the mean, such that the Fano factor is one. However, the Poisson model and thereby the FF do not apply to LFP power. To nevertheless arrive at a mean-adjusted metric of LFP power variability, we considered the coefficient of variation (CV), that is, the standard deviation (SD) divided by the mean. In order to properly calculate the CV of LFP power, we needed to make sure that the SD and the mean of LFP power were properly estimated, which required that LFP power values were approximately normally distributed. Raw LFP gamma power values in our dataset were highly non-normally distributed (Figure 5c, one-sample Kolmogorov-Smirnov test: p=3×10^−61^). Such LFP power spectra are typically normalized, e.g., by division by an appropriate control spectrum, like e.g. the LFP power spectrum during a baseline period or a control condition, or the 1/f spectrum fitted to the LFP power spectrum of interest. Here, we used a normalization by the pre-stimulus baseline. Because our LFP power ratio values were still highly non-normally distributed (Figure 5d, one-sample Kolmogorov-Smirnov test: p=1×10^−60^), we adopted best practices to log transform the power ratio values, and indeed, the log10 of the power ratios in our dataset were approximately normally distributed (Figure 5e, one-sample Kolmogorov-Smirnov test: p=0.232). This allowed the calculation of a proper mean (Figure 5f), SD (Figure 5g) and CV (Figure 5h). Intriguingly, the CV in the gamma band showed a pronounced reduction after stimulus onset, with a phasic and a sustained component (Figure 5i). Thus, stimulus onset quenched the variability of neuronal gamma-band activity in awake macaque area V1. In this particular dataset, visual stimulation induced not only a sustained gamma-power but also a sustained alpha-beta power enhancement (Figure 5j). Intriguingly, this enhancement in alpha-beta power was also accompanied by a decrease in the CV (Figure 5j).

Inspired by this finding, we last investigated whether similar effects were present in the ECoG dataset (Figure 6a) that we had already used for the ATV and ITV analysis above. Also in this dataset, gamma power values were highly non-normally distributed (Figure 6b, one-sample Kolmogorov-Smirnov test: p=3×10^−20^), and gamma power ratio values were highly non-normally distributed (Figure 6c, one-sample Kolmogorov-Smirnov test: p=4×10^−20^). As for the other dataset, the log10-transformed LFP gamma power ratios were approximately normally distributed (Figure 6d, one-sample Kolmogorov-Smirnov test: p=0.275). This allowed the calculation of a proper mean (Figure 6e), SD (Figure 6f) and CV (Figure 6g). Stimulus onset led to a similar reduction in gamma CV (Figure 6h) as in the microelectrode data (Figure 5i). Importantly, in this dataset, visual stimulation did not increase alpha-beta power, but it induced the classical alpha-beta power reduction (Figure 6b). Intriguingly, this alpha-beta power reduction was accompanied by an increase in the CV(log(alpha power ratio)) (Figure 6i).

**Figure 6.**
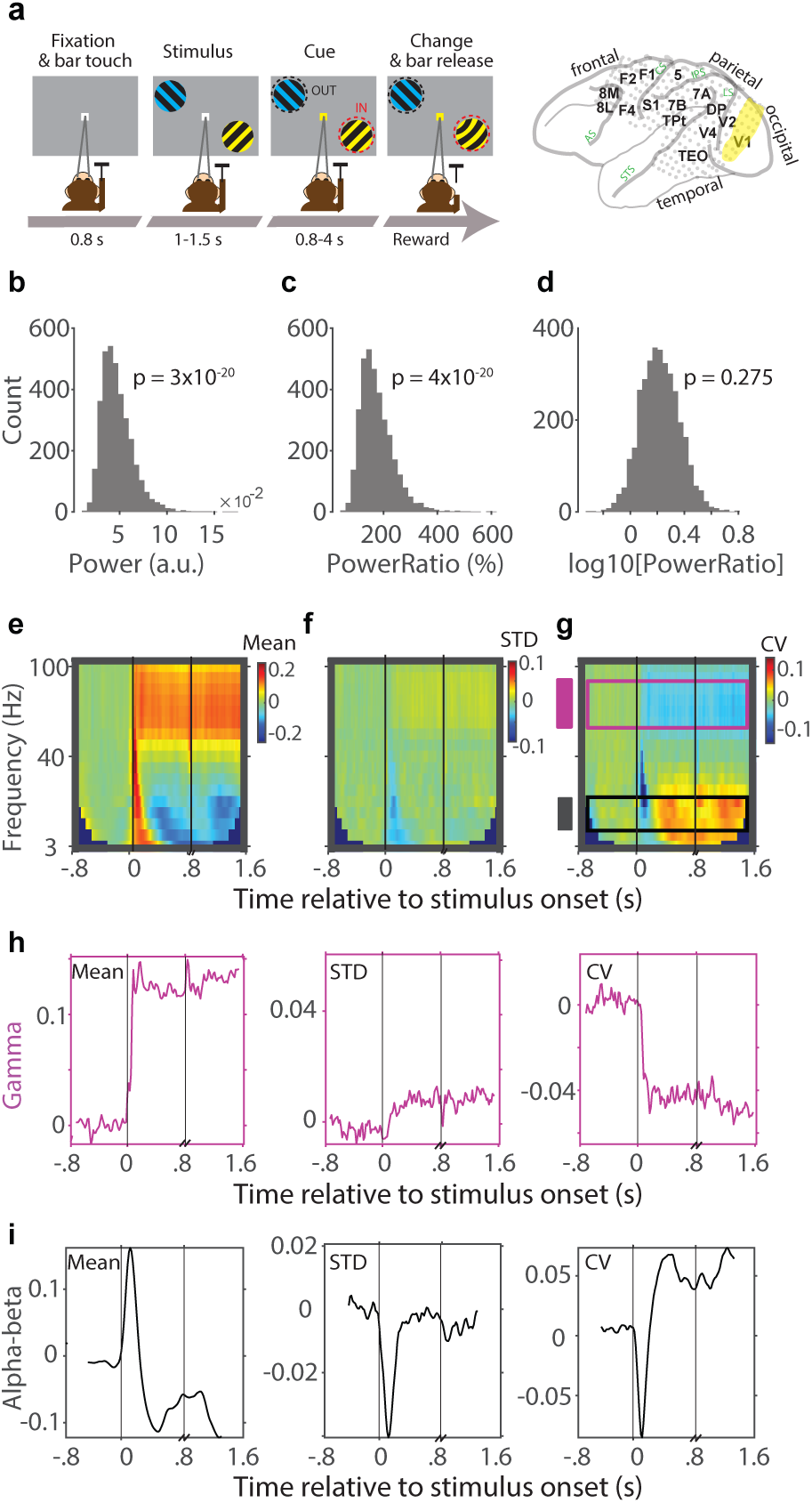
The Coefficient of Variation (CV) of log(power) as a mean-adjusted variance metric for LFP power. Data from subdural ECoG array. **a**) Schematic illustration of the attention task during which ECoG data were recorded. Data were recorded simultaneously from 252 electrodes distributed across the 15 indicated brain areas. The analysis used the data from area V1, highlighted in yellow. *AS*: Arcuate Sulcus, *CS*: Central Sulcus, *STS*: Superior Temporal Sulcus, *IPS*: Intraparietal Sulcus, *LS*: Lateral Sulcus. **b**) Distribution of raw LFP gamma-band power across sessions, channels and trials. **c**) Same as (**b**), but for the ratio of gamma-band power during stimulation over gamma-band power during pre-stimulus baseline. **d**) Same as (**c**), but taking the log10 of the ratio. All subsequent panels use this log10-ratio. **e-g**) Time-frequency representation of the log10-ratio mean (**e**), standard deviation (**f**), and coefficient of variation (**g**). Shown are the averages over 317 attention-IN trials. **h**) Cross-cuts for the gamma band indicated in (**g**) as purple rectangle and shown here as time-course to better illustrate the stimulus-related dynamics. **i**) Same as (**h**), but for the alpha band indicated in (**g**) as black rectangle.

## Discussion

We found that ATV is essentially identical to ITV for periods devoid of ERPs. The extremely high ATV-ITV correlations were revealed by calculating the two metrics in strictly corresponding ways, by first filtering the data and then computing variance either across trials (ATV) or within trials (ITV). Importantly, ITV is equivalent to the commonly used spectral power. Thus, the strong correlation between ITV and ATV suggests that ATV reflects spectral power during periods devoid of ERPs. For periods with ERPs, ATV is reduced compared to ITV. The ratio of ATV over ITV is strongly negatively correlated with the ratio of evoked power over total power, a metric of the ERP’s contribution to the LFP. This held particularly during periods when the ERP was strong. These effects were recapitulated in very simple simulations. We then presented a metric for the mean-adjusted variability for LFP power, namely the CV of the log10-transformed LFP power ratio. This metric decreased when gamma power increased during visual stimulation, and it showed an inverse relation to alpha-beta power, when alpha-beta power increased or decreased during visual stimulation, respectively.

EEG/LFP signals are fundamentally different from neuronal SCs. While SCs reflect the output of neurons along their axons, EEG/LFP signals reflect the synaptic inputs on their dendrites and somata and the corresponding post-synaptic currents (PSCs). The PSCs summate to the EEG/LFP when they are synchronized, and this synchronization is greatly aided by neuronal oscillations. Oscillations render the spike outputs of the respective neurons and the resulting PSCs at their target neurons partly temporally predictable, and thereby provide a natural way for oscillatory synchronization. Oscillatory synchronization occurs at characteristic frequency bands, and oscillations of various frequencies overlap to form the LFP, EEG and MEG. These oscillations have been linked to neuronal communication and processing ^24, 25^.

Neuronal responses, and also ensuing behavioral responses, to a given input can be modulated dependent on the oscillation phase ^26^. However, crucial for the present study, the momentary absolute phase of an ongoing oscillation does not seem to be a code that can be directly compared to the neuronal SC code. Let us assume that a particular stimulus induces a pattern of SCs across a group of neurons. Then, by recording this pattern of SCs for a short window of time, one can decode the stimulus with high fidelity. Let us now assume that these neurons, because they are composed of reciprocally coupled excitatory and inhibitory neurons, engage in an ongoing oscillation. Then, by recording and estimating the instantaneous oscillation phase, one can probably not decode the stimulus. The phase undergoes a full rotation during each oscillation cycle, such that it does not differentiate between stimuli as long as all stimuli induce the oscillation to some degree. Note that the stimulus might well be decoded from the pattern of phase relations between recording sites ^27, 28^ or from the pattern of oscillation strengths ^29–31^. This is different from the instantaneous oscillation phase of one particular recording site. Correspondingly, the phase of an ongoing neuronal oscillation at a particular time point typically varies randomly across trials, except when the oscillation is transiently reset, e.g., by a stimulus onset or a (micro)saccade. These resets can lead to ERPs ^23^, during which the phase and the corresponding amplitude of the LFP signal do carry information about the respective event. Yet, neuronal oscillations are not sinusoidal oscillations with linear phase evolution, but rather their phases drift substantially (which also contributes to them typically occupying a spectral band rather than a spectral line). Thereby, after a resetting event, the oscillation phase drifts and after few cycles is again random across trials. As the oscillation phase directly maps onto the raw instantaneous signal amplitude, this instantaneous amplitude value is equally random across trials. From these considerations, we suggest that the instantaneous amplitude of the ongoing LFP, EEG or MEG signal, outside of ERPs, is not suited for stimulus decoding in the same way as neuronal SCs. Correspondingly, while the ATV of SCs is highly meaningful, the meaning of the ATV of the instantaneous EEG amplitude is less clear.

Here, we have shown empirically that the ATV of the LFP (or EEG) amplitude is very highly correlated to the ITV of LFP (or EEG), that is to power, a standard metric derived from the LFP, EEG or MEG. This identity holds for periods outside of ERPs. During ERPs, the strength of the ERP determines how much ATV is reduced relative to the ITV. Because stimulation does induce reliable changes in LFP power, we investigated whether the variability of LFP power – rather than the raw LFP amplitude – showed interesting effects of stimulation. Indeed, we found that the CV of the logarithm of the LFP power ratio is a suitable metric. This metric, at least under simple circumstances, corresponds to the reciprocal of the standard d’ metric from signal detection theory, applied to the strength of the stimulus-elicited power Ps normalized by baseline power Pb. Note that if Ps and Pb are both lognormally distributed, then the log power ratio, log(Ps/Pb), is the difference of two Gaussian variables and its CV is the SD of the difference divided by the mean of the difference, i.e., the reciprocal of d’. From this perspective, when the CV measure becomes smaller after stimulus onset, the d’ based on the log-power-ratio metric increases. Intriguingly, this metric is constructed similarly to the FF of single-unit SCs, and it shows similar effects: For LFP gamma power, which is enhanced by stimulus onset, it is reduced by stimulus onset. For single unit SCs, FF reduction was established irrespective of whether stimulus onset resulted in a SC increase ^1^, while SC decreases were not studied. We had the opportunity to investigate LFP power decreases. In LFP recordings from ECoG, we found the classically described stimulus induced alpha-beta power decrease. Intriguingly, when stimulation decreased alpha-beta power, it increased the CV of log(alpha-beta power ratio). In a different dataset, recorded with intracortical electrodes, stimulation induced alpha-beta power increases, and decreases of the CV of log(alpha-beta power ratio). These findings suggest that changes in the CV of log(power ratio) might generally be negatively correlated to the corresponding changes in power in the same frequency bands. Yet, this needs to be tested across many datasets to arrive at a firm conclusion ^15^.

To our knowledge, the CV(log(power ratio)) has so far not been systematically studied as a function of stimuli and tasks ^32, 33^. We envisage that many such studies can easily be performed on existing and future LFP, EEG, and MEG datasets. These studies would thereby investigate whether and how these continuous neuronal signals show similar variability quenching as has been prominently described for single-unit SCs ^1^.

## Acknowledgements

This work was supported by DFG (FOR 1847 FR2557/2-1, FR2557/5-1-CORNET, FR2557/7-1-DualStreams), EU (HEALTH-F2-2008-200728-BrainSynch, FP7-604102-HBP), a European Young Investigator Award, and the National Institutes of Health (1U54MH091657-WU-Minn-Consortium-HCP, R00NS115918).

## Data availability

Data and code used for analysis in this study are available upon reasonable request.

## Lead contact

For further information or resource requests, please contact the Lead Contact, Mohsen Parto-Dezfouli.

## Author contributions

M.P. and P.F. conceptualized the study. C.A.B., E.L.J., and E.P. collected the ECoG, EEG, and Microelectrode data. M.P. and P.F. conducted the analyses and prepared the original draft. M.P., E.L.J., E.P., C.A.B., S.K., and P.F. jointly prepared the final manuscript.

## Supplementary information

Document S1. Figures S1-S2.

## Materials and Methods

We calculated ATV and ITV on an ECoG dataset from one awake macaque, on a microelectrode dataset from one awake macaque, an EEG dataset from human subjects, and on simulated data. Additionally, we quantified SC variability using the Fano Factor for the microelectrode dataset, and we quantified LFP power variability by the coefficient of variation for the microelectrode and the ECoG dataset. All empirical datasets have been acquired under respective ethical approvals. For the ECoG dataset, experimental procedures were approved by the animal ethics committee of Radboud University Nijmegen (Nijmegen, the Netherlands). For the intracortical microelectrode dataset, experimental procedures were approved by the regional authority (Regierungspräsidium Darmstadt, Germany). For the human EEG data, written informed consent was obtained in accordance with the University of California, Berkeley, Institutional Review Board or the Regional Committee for Medical Research Ethics. All datasets have been used in previous publications, which describe the experiments in detail ^21, 22, 34-38^. The following methods description will focus on the specific data analyses for the current study.

### Datasets

#### ECoG dataset

In the ECoG dataset, local field potentials were recorded using an ECoG electrode grid implanted on the left hemisphere of two macaque monkeys (Monkey K and Monkey P) ^35^. For the current study, we used data from one monkey (Monkey K) ^39^. The monkey was trained to perform a selective visual attention task, requiring it to maintain its gaze on a central fixation point and pressing a lever throughout a trial. After an 0.8 s fixation period, two patches of sinusoidal grating appeared, drifting behind a circular aperture, one colored blue, the other yellow, with the colors randomly distributed across trials. A subsequent change in the fixation point’s color cued the monkey to attend to the stimulus with the matching color. The monkey had to release the lever in response to a shape change in the attended stimulus to obtain a reward, while ignoring changes in the non-attended stimulus. The monkey achieved 94% accuracy. During the task, LFPs were recorded from 252 channels across 15 brain areas. We have previously established that the two most prominent brain rhythms in these data are the gamma rhythm in the occipital region (defined here as areas V1, V2, V4) and the beta rhythm in the frontal region (defined here as areas F1, F2, F4) ^21^. Correspondingly, the current analysis focused on those rhythms and regions. Note that the frequency-band definitions in the present study differed slightly from those in the previous study, and we here used a combined alpha-beta band. Trials with attention directed to the visual hemifield contralateral to the recorded hemisphere were referred to as attend-IN (or just IN) condition, and trials with attention ipsilateral as attend-OUT (or just OUT) condition. We considered only trials with at least 0.8 s stimulus presentation before the cue and 0.8 s between the cue and the stimulus change.

For Figures 1, 2 and 6, we segmented the data as follows: (1) “fixation” period: last 0.8 s of fixation end-aligned to stimulus onset; (2) “stimulus” period: first 0.8 s after stimulus onset; (3) “cue” period: first 0.8 s after cue onset; (4) “change” period: last 0.6 s end-aligned to stimulus change. For Figure 3, we segmented the data differently, because we aimed at contrasting periods with and without ERP: (1) “fixation” period: last 0.5 s of fixation end-aligned to stimulus onset; (2) “ERP” period: first 0.3 s after stimulus onset; (3) “sustained” period: the 0.5 s between 0.3 and 0.8 s after stimulus onset.

#### EEG data

The EEG dataset was collected from two groups of participants: 14 individuals with unilateral prefrontal cortex (PFC) lesions (lesion group) and 20 age- and education-matched healthy controls (control group). For the current study, we analyzed the EEG data from one human participant from the control group. Participants completed a working memory (WM) task consisting of four phases: (1) baseline, (2) encoding, (3) maintenance, and (4) active processing. Each trial began with a 2-second fixation, followed by two sequential stimuli (0.2 s each) with a 0.2 s inter-stimulus interval. After a brief blank period (duration randomly either 0.9 or 1.15 s), participants were asked to identify the relation (TOP/BOTTOM/FIRST/SECOND) or identity (SAME/DIFFERENT) between the two previously presented stimuli, followed by another delay period before responding by selecting the correct stimulus. EEG signals were recorded from 64+8 channels using a BioSemi ActiveTwo amplifier, with Ag-AgCl electrodes positioned according to the 10-20 system ^22^.

#### Microelectrode data

LFP and single-unit spike data were intracortically recorded from areas V1 and V2 of a macaque monkey during a fixation task. The monkey maintained central fixation throughout each successfully completed trial and was rewarded for this with grape juice. Following a 1-1.1 s baseline period, the monkey was presented with a grating stimulus for 1-1.3 s. Grating orientations varied from 0° to 170° in 10° steps, with one orientation presented per trial. Each recording session consisted of three stimulus blocks. In the first and third block, all orientations were presented in a pseudorandom sequence. In the second block, one specific orientation was randomly selected for each recording session and was presented 90 times in a row. We analyzed the 90 consecutive trials of the same stimulus from the second block. Spike data were sorted into single units using SpyKING CIRCUS ^40^.

#### Simulated data

The simulated data were drawn from two distributions, representing a fixation period (0.5 s) and a stimulus period (1 s). The distributions were standard normal distributions according to the following formula:

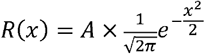

where *x* is a random variable with a mean of 0 and variance of 1, and *A* represents the standard deviation of the distribution.

### Analysis

All analyses were performed using MATLAB 2020b utilizing the FieldTrip toolbox ^41^. Line-noise artifacts were removed for the ECoG data at 50 Hz and harmonics and for the EEG data at 60 Hz and harmonics. For the microelectrode data, line-noise removal was not necessary. All LFP and EEG signals were normalized (z-transformed by subtraction of their mean and division by their SD) and detrended. ATV was estimated based on across-trial variance for each 1 ms time bin, and smoothed using a 20 ms sliding window. ATV was calculated as follows:

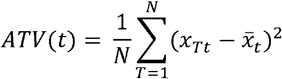

where *x*_*Tt*_ is the amplitude of the signal at time *t* in trial *T*, 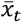 is the mean amplitude over trials at time *t*, and N is the number of trials.

ITV was calculated as follows:

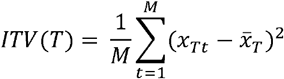

where *x*_*Tt*_ is the amplitude of the signal at time *t* of trial *T*, 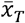 is the mean amplitude within trial *T*, and M is the number of time points.

In the ECoG dataset, both ATV and ITV were calculated separately for each of 15 brain areas, and the Pearson regression coefficient between ATV and ITV across brain areas was assessed for different periods as described above and in the Results section.

For power-spectrum analysis, the signals (of the respective period lengths, as described above and in the Results section) were Hann-tapered, zero-padded to 1 s, and Fourier transformed. Power spectra were computed as the squared magnitude of the Fourier transform.

To test for the normality of the distributions of power, power ratio, or logarithm of power ratio, we used the One-sample Kolmogorov-Smirnov test (MATLAB function ‘*kstest*’). It determines whether a distribution comes from a standard normal distribution or not. P-values greater than 0.05 indicate that the corresponding distribution is normal and vice versa.

