## Supplementary material for "On variability in local field potentials": Document S1. Figures S1-S2.

### Supplementary information

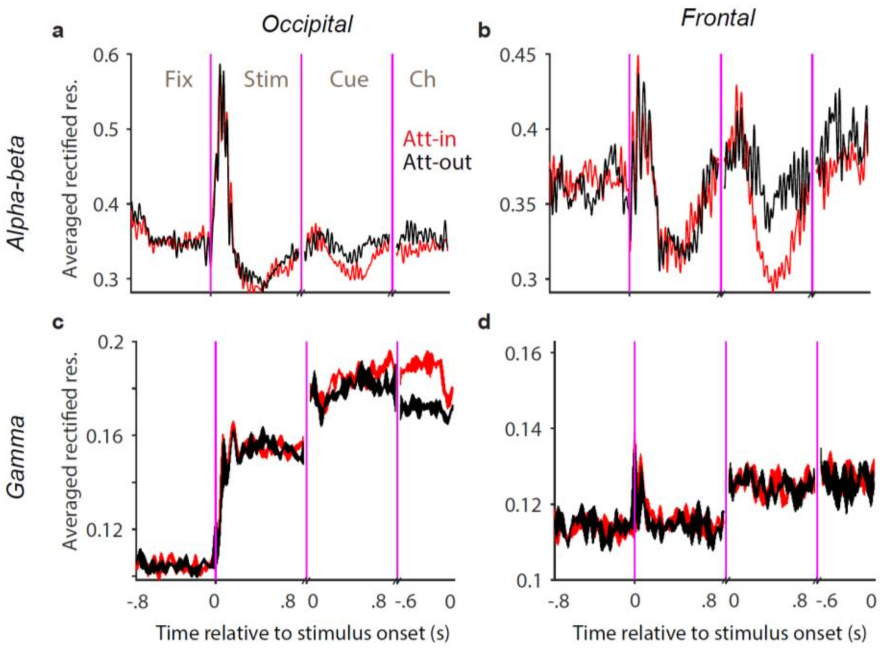


**Figure S1. The rectified LFP averaged over trials reflects ATV of LFP.**

**a, b**) The trial-averaged rectified LFP, after filtering in the alpha-beta band (8–18 Hz) for attend-IN (red) and attend-OUT (black) conditions, shown for the occipital (**a**) and frontal (**b**) cluster.
**c, d**) Same as (**a, b**), but for data filtered in the gamma band (70–80 Hz).

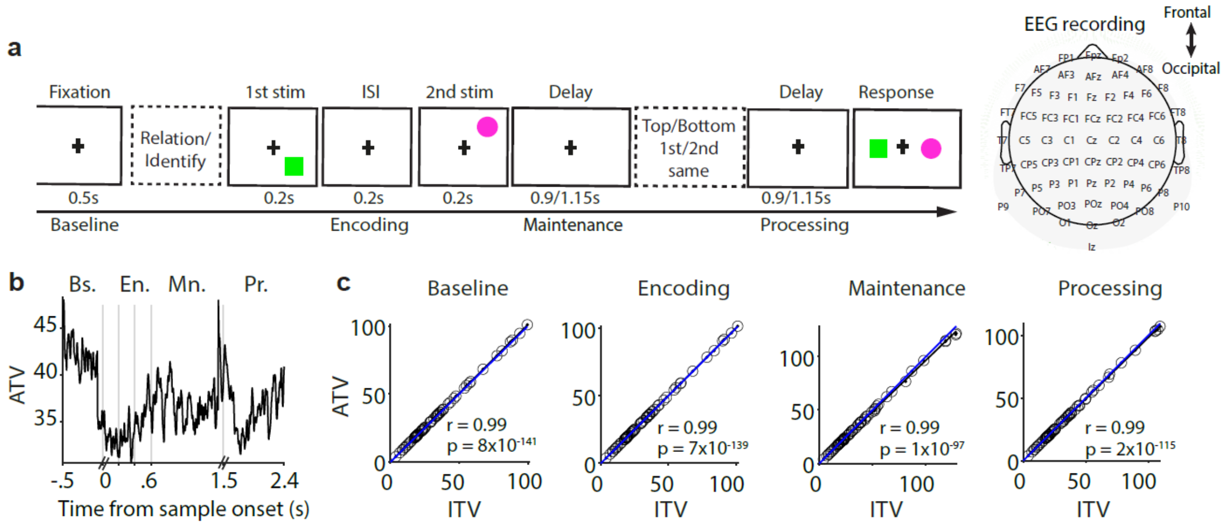


**Figure S2. ATV correlates with ITV for EEG data.**

**a**) Schematic of the behavioral working memory (WM) task consisting of four task periods: baseline, encoding, maintenance, and active processing. Subjects maintained the spatial and temporal features of stimuli while performing the WM task. Data were recorded from 64 EEG electrodes.

**b**) Time-course of ATV during the four task periods for one example subject averaged across 64 recording channels. *Bs*: baseline, *En*: encoding, *Mn*: maintenance, and *Pr*: active processing.

**c**) Correlation between ATV and ITV during the four task periods. Each dot represents data from one recording channel.
